# Fiber-tract development contributes to functional specialization in the human hippocampus

**DOI:** 10.64898/2026.04.28.721442

**Authors:** Jonah Kember, Christine L. Tardif, Sylvain Baillet, Ying He, Sam Audrain, Alex Barnett, Tracy Riggins, Xiaoqian J. Chai

## Abstract

Fiber-tracts exhibit distinct projection patterns along the anterior–posterior axis of the hippocampus, promoting a specialization in function. This specialization becomes increasingly pronounced throughout child development, with important implications for neurocognitive outcomes. Developmental changes in fiber-tract properties, including intra-axonal cross-sectional area and myelin content, may contribute to this anterior-posterior functional specialization. To test this, we developed a diffusion-MRI tractography pipeline to identify hippocampal fiber-tracts in single subjects, then examined whether age-related differences in total intra-axonal cross-sectional area and myelin content (T1w/T2w) could predict functional specialization in a large cross-sectional sample (*N*=539, aged 5–21 years). With age, we found that the cross-sectional area of short-range medial-temporal tracts, which primarily target the anterior/body of the hippocampus, exhibited rapid growth. Concomitantly, the cross-sectional area of long-range occipito-parietal tracts, which primarily target the posterior hippocampus, exhibited a modest pruning. Increases in myelin content were relatively homogenous across fiber-tracts. In support of our hypothesis, we found that the cross-sectional area of fiber-tracts significantly predicts the surface-area of an fMRI-defined posterior system; a sensitive marker of functional specialization in the hippocampus. Tracts targeting early visual cortex (V2, V3, V4) showed the strongest association, with statistical modeling indicating a mediating effect of early-visual tract development on the relation between age and functional specialization. These findings provide evidence consistent with a mechanism whereby anatomical neurodevelopment contributes to functional specialization in the human hippocampus.

## Main

In the human hippocampus, distinct gene-expression domains along the longitudinal axis give rise to anterior–posterior differences in the projection patterns of fiber-tracts (i.e., bundles of myelinated axons; Ayhan et al., 2021; Strange et al., 2014; Thompson et al., 2008; Vogel et al., 2020). In the posterior hippocampus, fiber-tracts preferentially project to medial occipital- and parietal-cortices; in the anterior hippocampus, fiber-tracts preferentially project to medial-temporal and subcortical regions (e.g., amygdala, thalamus; Dalton et al., 2022; Huang et al., 2021; Insausti & Muñoz, 2021). These projection patterns promote an anterior–posterior specialization of function, with the posterior hippocampus showing increased sensitivity to visuo-spatial information (Nichols et al., 2023). Indeed, in rodents, lesioning the dorsal (corresponding to the posterior hippocampus in humans), but not ventral hippocampus (anterior in humans) severely impairs spatial memory, and dorsal neurons are approximately ten times more sensitive to spatial information than ventral neurons (place-field widths of ∼1m vs ∼10m; Pothuizen et al., 2004; Kjestrup et al., 2008; Strange et al., 2014).

This specialization of function is reflected in partially dissociable hemodynamic signaling between anterior and posterior regions, and can therefore be empirically observed with functional-MRI (Angeli et al., 2025; Audrain et al., 2024; Nichols et al., 2025; Xie et al., 2024; Zheng et al., 2021). Precision-mapping techniques, which use extended acquisition time fMRI to assign individual patches of hippocampal tissue to anterior- and posterior-functional systems, have proven particularly sensitive to this specialization at the single-subject level (Angeli et al., 2025; Kember et al., 2026; Zheng et al., 2021). Using this approach in a cross-sectional developmental sample, we recently estimated that the surface-area of hippocampal tissue dedicated to the posterior functional system decreases at a rate of approximately –21.4 mm^2^ per year (95% CI: −25.3, −17.6) from age 5 until at least late adolescence (Kember et al., 2026). The surface-area of the anterior functional system increases at a similar albeit slower rate (16.2 mm^2^ / year [95% CI: 11.5, 20.8]). This is in line with findings indicating a posterior-to-anterior shift during development of the principal functional-gradient of the hippocampus, which extends along the longitudinal axis (Xie et al., 2024).

The functional development of the hippocampus has substantial clinical relevance. Hippocampal dysfunction is associated with deficits in learning, visuo-spatial memory capacity, and socioemotional regulation (Barch et al., 2020; Gilchrist et al., 2018; Kember et al., 2026; Tang et al., 2020; Xie et al., 2024); and underlies the presence of these deficits across a range of neurodevelopmental conditions, including those linked to early developmental insult (e.g., hypoxia, placental insufficiency, fetal alcohol exposure) and genetic risk (e.g., autism; Kember et al., 2024; Li et al., 2019; White et al., 2024).

Despite its importance, the mechanisms driving increased functional specialization along the anterior–posterior axis during human development have not yet been identified. In general, the mechanisms underlying hemodynamic differentiation in the hippocampus are well understood: cortico-hippocampal fiber-tracts deliver different synaptic inputs along the anterior-posterior axis, which in turn bias the targeted hippocampal areas towards distinct patterns of neuronal, and consequently hemodynamic, activity (Almeida & Lyons, 2018). With this mechanism in mind, we hypothesized that developmental increases in functional specialization might be driven by the development of these cortico-hippocampal fiber-tracts.

To non-invasively test for evidence of this mechanism, we mapped the precise pathway of hippocampal fiber-tracts in single subjects and quantified their strength through diffusion-MRI tractography. To quantify fiber-tract strength, we focused on properties thought to promote the efficacy of synaptic transmission, namely: tract size (fixel-based intra-axonal cross-sectional area, measured in mm^2^; Sotiropoulos & Zalesky, 2017; Smith et al., 2015; Smith et al., 2022), and myelin content (the ratio of T1-weighted to T2-weighted anatomical contrasts; Glasser & Van Essen, 2011). We then applied this technique in a large cross-sectional sample (*N*=539; aged 5-21) in which high-resolution diffusion-MRI and extended acquisition-time functional-MRI had been acquired, then tested whether differences in fiber-tract cross-sectional area and myelin content could predict the surface-area of functional systems along the longitudinal axis. In doing so, we found correlational evidence in support of our hypothesized mechanism. Specifically, that functional specialization in the human hippocampus is partially attributable to the pruning of long-range fiber-tracts between the posterior hippocampus and occipital cortex. These findings are well supported by prior work, and provide initial evidence that structural changes to hippocampal fiber-tracts contribute to the development of anterior–posterior functional specialization in the hippocampus.

## Results

### Quantifying hippocampal white-matter tracts through diffusion-MRI tractography

In each subject, the gray-matter/white-matter interface of the hippocampus was identified from a volumetric segmentation of the hippocampus, which was obtained from the 0.8mm resolution T1-weighted image through *HippUnfold* (DeKraker et al., 2020). Fiber orientation densities (FODs) were then estimated using the multishell diffusion-MRI data (1.5mm voxels; 185 directions at b=1500 s/mm^2^ and 3000 s/mm^2^, 28 b=0 s/mm^2^ images), and were intensity normalized to allow for quantitative comparison of streamlines across subjects (Raffelt et al., 2017). Using these hippocampal masks and fiber orientation densities, we generated 1 million anatomically constrained streamlines; five hundred thousand seeded from each hippocampus (relevant parameters: min-length: 3.0 mm; max-length: 150.0 mm; algorithm: iFOD2; - crop_at_gmwmi; -backtrack; -seed_unidirectional). SIFT2 filtering of streamlines was then conducted to derive biologically accurate, quantitative estimates of cross-sectional area (in units mm^2^; Calamante et al., 2019; Dalton et al., 2022; Smith et al., 2015). Further processing details are provided in the methods.

On average across subjects (*N*=539, uniformly sampled from ages 5 to 21; 289 female, 250 male), the mean cross-sectional area of all hippocampal streamlines was 656.9 ± 235 mm^2^ combined across hemispheres. Of this cross-sectional area, 82.2% projected to medial-temporal lobe (MTL) regions, 10.82% projected to non-MTL cortical regions [Posterior_Cingulate: 16.98 mm^2^; Ventral_Stream_Visual: 15.95 mm^2^; Early_Visual: 8.89 mm^2^; Lateral_Temporal: 7.13 mm^2^; Primary_Visual: 6.21 mm^2^; Dorsal_Stream_Visual: 4.54 mm^2^; MT+_Complex_and_Neighboring_Visual_Areas: 2.04 mm^2^; all others < 2mm^2^], and 6.98% projected to subcortical regions [Amygdala: 22.14 mm^2^; Caud: 9.29 mm^2^; PuM: 9.17 mm^2^; all others < 2mm^2^].

We next grouped these streamlines into fiber-tracts (sets of streamlines with distinct hippocampal and cortical endpoints) using a custom template-matching approach relying on the QuickBundles algorithm (Garyfallidis et al., 2012). We identified 43 fiber-tracts projecting to the: Posterior_Cingulate cortex (11 tracts; 9.48 ± 3.32 mm^2^), Early_Visual cortex (11 tracts 10.91 ± 4.35 mm^2^), Dorsal_Stream_Visual cortex (7 tracts; 5.49 ± 2.91 mm^2^), Primary_Visual cortex (5 tracts; 3.60 ± 2.52 mm^2^), Lateral_Temporal cortex (5 tracts; 3.43 ± 2.55 mm^2^), Ventral_Stream_Visual cortex (3 tracts; 3.12 ± 1.75 mm^2^) and Superior_Parietal cortex (1 tract; 0.43 ± 0.37 mm^2^).

### Long-range occipital-parietal streamlines primarily target the posterior hippocampus

We next examined how the fiber-tract projection patterns differ between the anterior, body, and posterior hippocampus (see methods for details on long-axis ROIs). In line with prior work, we found that the posterior hippocampus has a considerably higher amount of cross-sectional area concentrated in long-range projections when compared to the body and anterior hippocampus (see Fig. 1A-C). Indeed, on average across subjects, the posterior hippocampus has 42.9 ± 15.2 mm^2^ of cross-sectional area in streamlines longer than 40 mm in length; while the body has 11.2 ± 5.8 mm^2^; and the anterior hippocampus has only 2.6 ± 3.0 mm^2^. This contrast between anterior and posterior hippocampal projection patterns was also evident upon visual inspection of single-subject tractograms (Fig. 1C).

**Fig 1.**
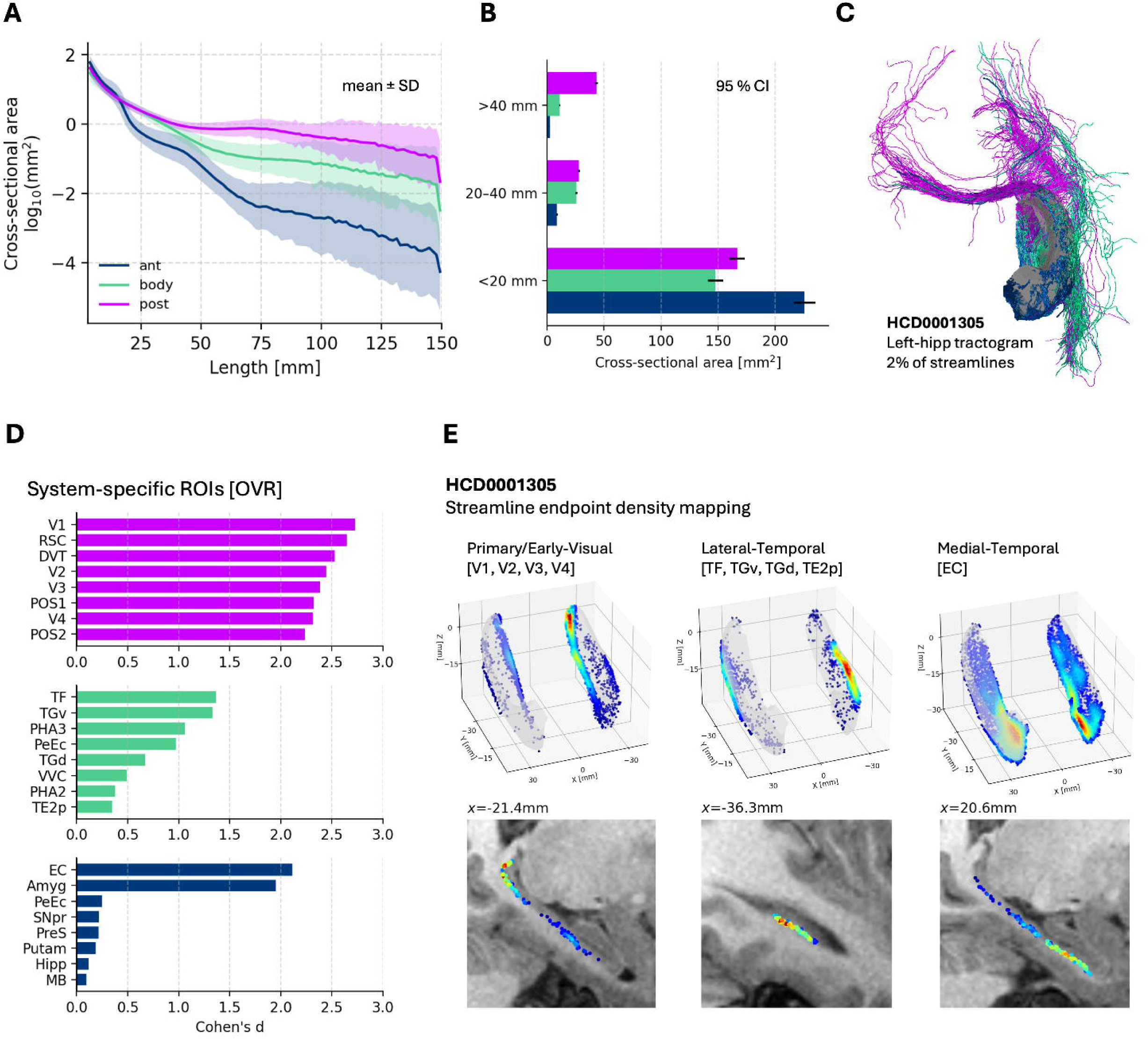
Long-range occipital-parietal streamlines primarily target the posterior hippocampus. (A) The mean ± SD cross-sectional area of hippocampal streamlines belonging to each hippocampal system (anterior, body, posterior) as a function of streamline length. Areas are log_10_ transformed to facilitate visualization. (B) Differences in the average cross-sectional area of streamlines belonging to each hippocampal system, for streamlines of varying length (short: < 20m, medium: 20-40mm inclusive, long: >40mm). Error bars reflect 95% confidence intervals. (C) Example single-subject tractogram (11.9 year old male), with streamlines colored by their termination in the hippocampus (pink: posterior, green: body, blue: anterior). (D) One-versus-rest analysis showing the ROIs which preferentially terminate in a specific hippocampal system. (E) For a single example subject, the density of streamline endpoints are plotted. On the top: endpoint density is overlaid on a 3D hippocampal surface. On the bottom: endpoint density is overlaid on a sagittal slice of the T1-weighted image.

Next, we identified the cortical regions that preferentially connect to each system along the hippocampal long-axis using a one-versus-rest analysis. Specifically, for each Glasser ROI, we calculated the cross-sectional area of streamlines targeting one system (e.g., posterior) and compared it to the cross-sectional area of streamlines targeting the other two systems (e.g., anterior + body) using a measure of effect-size (Cohen’s *d*; Fig. 1D). We found that the posterior hippocampus has projections which preferentially terminate in the occipital-parietal cortex (top-8 ROIs with highest specificity for posterior hippocampus: V1, V2, V3, V4, RSC, POS1, POS2, DVT); the body of the hippocampus has shorter projections that preferentially terminate in the lateral temporal cortex (ROIs with highest specificity for the hippocampal body: TF, TGv, PHA3, PeEc, TGd, VVC, PHA2, TE2p); while the anterior hippocampus has even shorter connections that preferentially terminate in subcortical and medial-temporal areas (ROIs with highest specificity for the anterior hippocampus: EC, Amygdala, PeEc, SNpr, PreS, Putamen, Hipp, MB).

Overall, these findings demonstrated a striking contrast between the posterior and anterior hippocampus, whereby long-range occipital-parietal tracts primarily target the posterior hippocampus; and short-range medial-temporal tracts primarily target the anterior hippocampus. These findings replicate prior diffusion-tractography findings in humans (Dalton et al., 2022; Huang et al., 2021), and tract-tracing findings in Macaques (Insausti & Muñoz, 2021). Indeed, in line with work in adults (Dalton et al., 2022), we found when inspecting the distribution of streamline endpoints in the hippocampi of single-subjects that streamlines targeting early visual cortex terminated in posterior-medial hippocampus; streamlines targeting lateral-temporal cortex terminated in the lateral hippocampal-body; and streamlines targeting medial-temporal cortex terminated in the hippocampal head (see Fig. 1E).

### Long-range occipital-parietal tracts show slight pruning with age while short-range medial-temporal tracts show rapid growth

Next, we sought to understand how fiber-tract cross-sectional areas differ with age. Overall, we found that the mean cross-sectional area of all hippocampal streamlines showed a moderate-to-large effect-size increase with age (Pearson’s *r* = .42, *p* < 1e-25). This was true for each hippocampal system, although it was significantly accelerated in the anterior hippocampus relative to the body and posterior hippocampus, as shown in Fig. 2A (anterior: 11.1 mm^2^ / year [95% CI: 9.1, 13.2]; body: 6.7 mm^2^ / year [95% CI: 5.3, 8.2]; posterior: 6.1 mm^2^ / year [95% CI: 4.8, 7.5]). When examining the specific fiber-tracts that contribute to this growth, we found that short-range medial-temporal tracts increased at an estimated rate of 26.1 mm^2^ / year across the sampled age range (Fig. 2B). In contrast, long-range cortical tracts showed a slight decrease in cross-sectional area with age (Fig. 2C). This effect was largest in magnitude for posterior-cingulate [-.24 mm^2^ / year, *p* < 1e-10], early-visual [-.28 mm^2^ / year, *p* < 1e-10], dorsal-stream-visual [-.18 mm^2^ / year, *p* < 1e-3], and primary-visual [-.25 mm^2^ / year, *p* < 1e-10] cortex.

**Fig 2.**
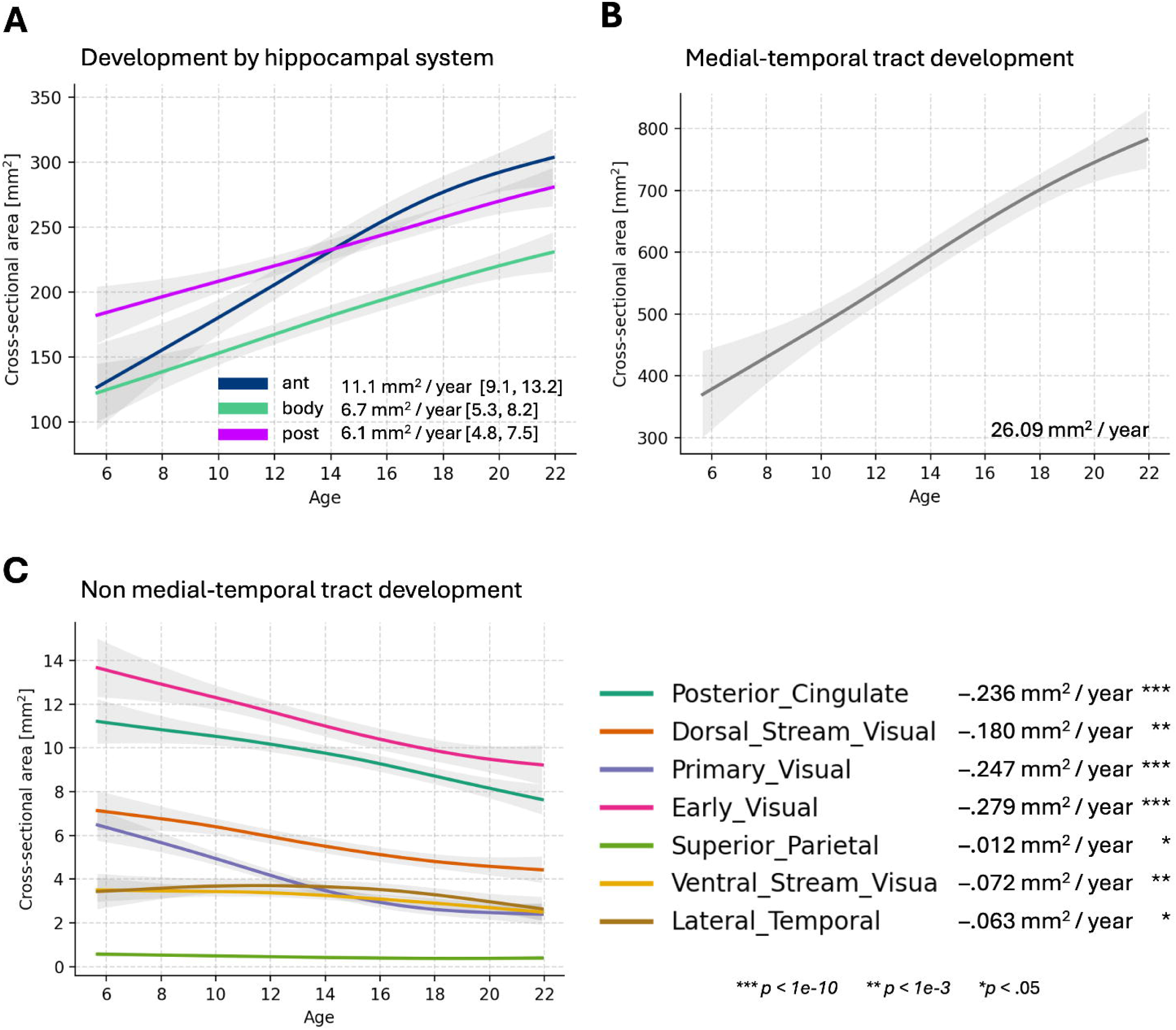
Long-range occipital-parietal tracts show slight pruning with age while short-range medial-temporal tracts show rapid growth. (A) Age-related differences in cross-sectional area of streamlines innervating the anterior, body, and posterior hippocampus with age. (B) Age-related differences in cross-sectional areas of fiber-tracts belonging to the medial-temporal cortex. (C) Age-related differences in cross-sectional areas of fiber-tracts belonging to different cortices. All parameters were estimated with linear-regression. Non-linear relations between cross-sectional area and age were estimated with a generalized additive model.

While small in magnitude when compared to the growth of medial-temporal fiber-tracts, these findings are indicative of a developmental pruning of the long-range hippocampal-cortical tracts targeting occipital-parietal cortex. It is important to note that fixel-based cross-sectional area reflects properties related to macrostructural white-matter bundles, and does not uniquely distinguish axonal pruning from changes in axon diameter, packing density, or extracellular space. Yet, this finding was unexpected, as prior work indicates an increase in the strength of long-range connections across the cortex (although not necessarily the hippocampus) during the same developmental period (Hagmann et al., 2010). Thus, to further confirm the veracity of this finding, and to understand the specificity of this effect to the hippocampus, we generated whole-brain tractograms using an identical tractography pipeline, and examined how the cross-sectional area of long-range projections change with development for each cortical ROI in the Glasser atlas. In contrast to the hippocampal effect, we found that the majority of cortical regions (116/180) actually showed an *increase* in the prominence of long-range fiber tracts with age (measured via the streamline length at 50% the cumulative sum of cross-sectional area; see supplementary Fig. 1), in line with prior work (Hagmann et al., 2010). We also noticed a common trend across the cortex, whereby regions which tend to have longer fiber-tracts also tend to show an increase in the concentration of fiber-tract area within long-range connections; and vice-versa for regions with shorter tracts. This helped us contextualize why medial-temporal ROIs, which tend to have very short-range connections relative to the rest of the cortex (a finding shown in our data as well as that of others; Bajada et al., 2019) show a developmental decrease in the area of long-range fiber-tracts. Thus, despite being initially unexpected, this gave us confidence in our finding that long-range occipital-parietal projections with the hippocampus show a slight pruning with age.

### Long-range posterior-hippocampal fiber tracts are heavily myelinated and undergo myelinogenesis

To estimate myelin content of hippocampal fiber-tracts, T1w/T2w values were sampled along each hippocampal streamline, and the median value was taken as a measure of streamline myelin content (Glasser & Van Essen, 2011). We note that although T1w/T2w does not constitute a quantitative or histologically specific measure of myelin content, it provides a myelin-sensitive contrast. For tract summary statistics, the median streamline T1w/T2w was weighted by streamline cross-sectional area (weights were normalized to maintain units). On average across subjects, we found that streamlines terminating in the posterior hippocampus had the highest T1w/T2w values (2.72 ± .26), followed by streamlines terminating in the body (2.34 ± .21), and streamlines terminating in the anterior hippocampus (1.92 ± .15). Each of these was a large-effect size difference (pairwise Cohen’s *d:* ant–body = 2.31; ant–post = 3.75; body–post = 1.59; *p*’s < .001; see Fig. 3A). This posterior–anterior gradient in T1w/T2w contrast is in line with the general principle observed across the cortex, where spatially proximal structural connections show reduced myelination, while long-range streamlines, primarily those innervating sensory cortices, show heavy myelination (Sandrone et al., 2023; Glasser et al., 2011). We also found that this general pattern could be appreciated through visual inspection of the subject-level T1w/T2w images, which we have highlighted in Fig. 3B.

**Fig 3.**
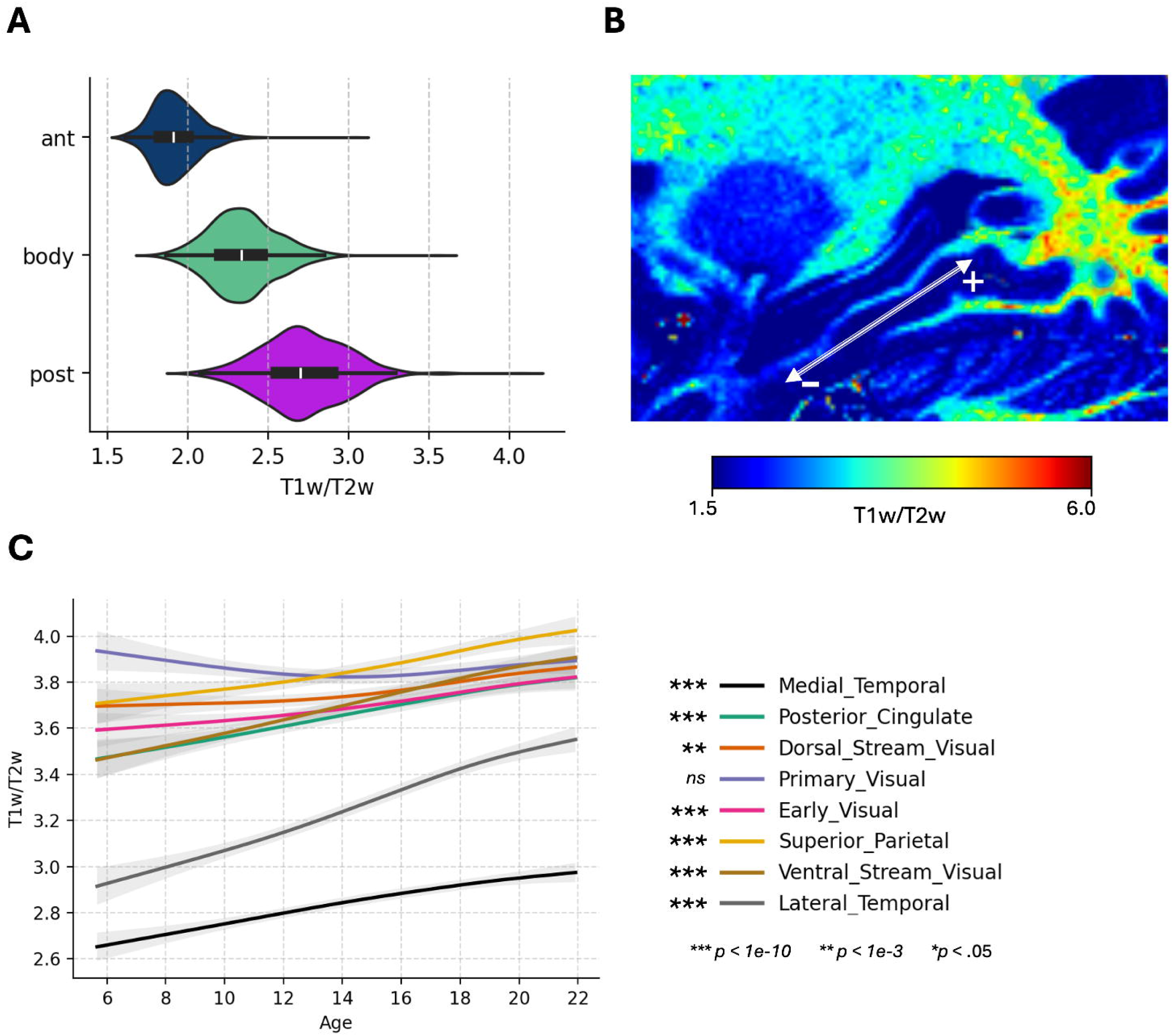
Long-range posterior-hippocampal fiber tracts are heavily myelinated and undergo myelinogenesis. (A) Differences in myelin content (T1w/T2w) for streamlines belonging to the anterior, body, and posterior hippocampal systems. (B) Sagittal slice of a T1w/T2w map for a single example subject, illustrating that the anterior-posterior gradient in myelin content for hippocampus-adjacent white-matter can be appreciated at the single-subject level. (C) Age-related differences in myelin content of fiber-tracts belonging to different cortices, along with parameter estimates (linear-regression predicting age from myelin content). Non-linear trajectories were estimated with a generalized additive model.

When inspecting fiber-tracts, we found that long-range occipital-parietal tracts had higher T1w/T2w compared to short-range medial-temporal and lateral-temporal tracts (see Fig. 3C). Tracts with the highest T1w/T2w contrast were those terminating in: primary_visual (3.87 ± .17), superior_parietal (3.86 ± .28), dorsal_stream_visual (3.73 ± .17), ventral_stream_visual (3.71 ± .12), early_visual (3.70 ± .20), and posterior_cingulate (3.67 ± .33) cortex; those terminating in lateral_temporal (3.26 ± .35) and medial_temporal (2.85 ± .75) cortex had relatively lower T1w/T2w values. All pairwise differences (i.e., between cortices) were significant (*p* < .05) except for primary_visual versus superior_parietal (*p* = .13).

With age, the majority of fiber-tracts showed increased T1w/T2w values (suggesting myelinogenesis; see Fig. 3C). Indeed, significant increases were observed for fiber-tracts terminating in medial_temporal, lateral_temporal, superior_parietal, early_visual, dorsal_stream_visual and posterior_cingulate cortex (all *p*-values < 1e-3). Fiber-tracts terminating in the primary_visual cortex, despite showing the highest absolute T1w/T2w values across subjects, did not show significant age-related differences (*p* > .05). Non-linear developmental patterns are shown in Fig. 3C.

Together these results indicate that long-range fiber-tracts projecting between posterior hippocampus and occipital-parietal cortex show the highest T1w/T2w contrast, and that increases in T1w/T2w contrast are relatively common across hippocampal fiber-tracts.

### The size of the posterior functional system is predicted by the cross-sectional area of hippocampal fiber tracts

We next tested whether individual differences in the cross-sectional area of hippocampal fiber tracts are related to individual differences in functional specialization of the hippocampus. To do so, we tested whether the surface-area of each functional system (anterior, body, posterior) could be significantly predicted by fiber-tract cross-sectional areas through ridge regression (i.e., linear regression with L2-regularization to reduce overfitting). To avoid over-estimating model performance, and to account for the fact that the number of tracts varied across cortices, performance was estimated through 5-fold cross-validation (the average out-of-sample *R^2^* across folds; values above zero indicate improved out-of-sample prediction relative to a null model predicting the sample mean). Model significance was assessed through non-parametric permutation testing (*N*=500 permutations). Results are presented in Fig. 4.

**Fig 4.**
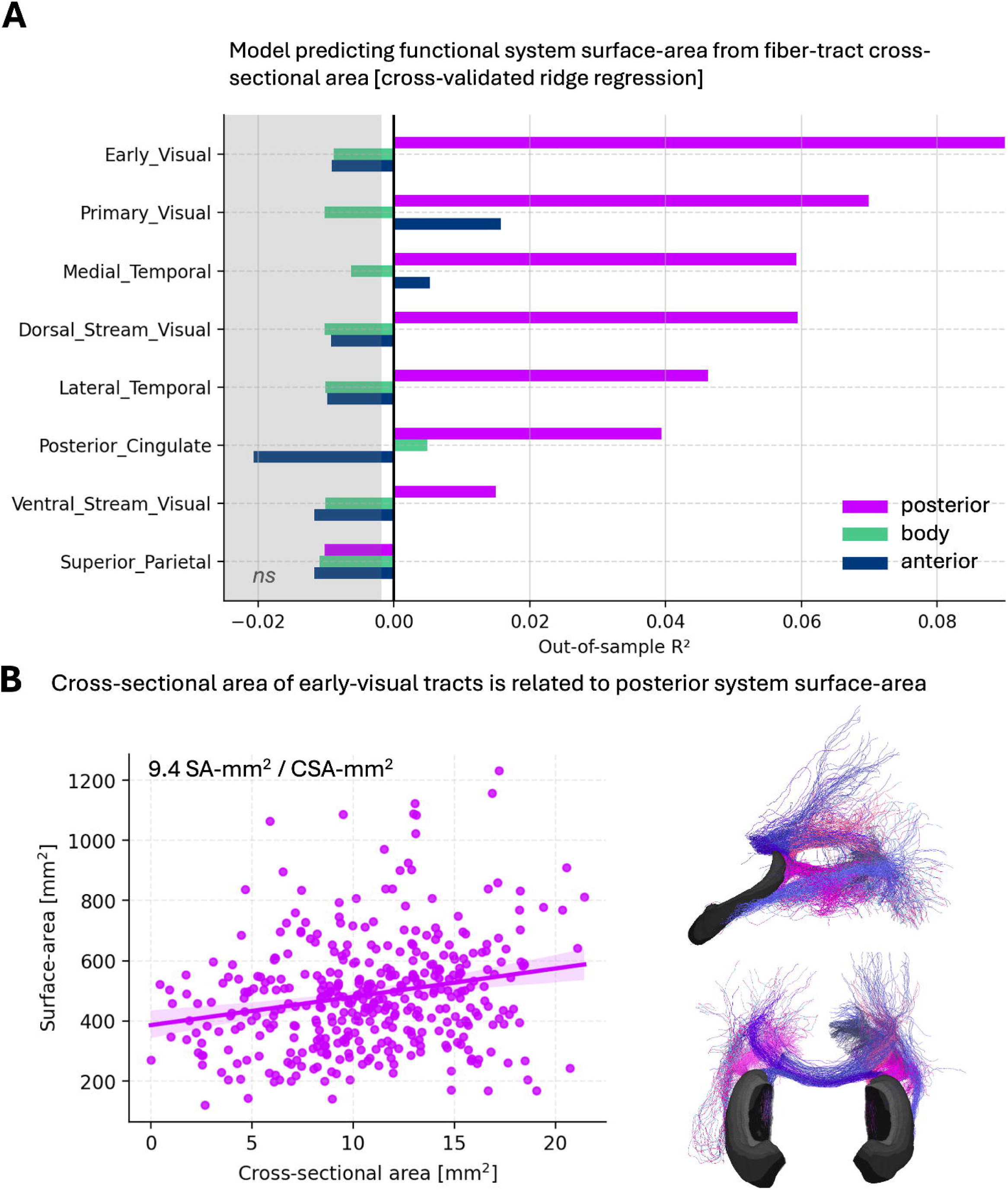
The size of the posterior functional system is predicted by the cross-sectional area of hippocampal fiber tracts. (A) Barplot indicating the out-of-sample performance (5-fold cross-validation) of ridge-regression models trained to predict functional-system surface-area (separately for: anterior, body, posterior) from the cross-sectional area of fiber-tracts (separately for tracts belonging to different cortices) across subjects. Values in the gray indicate model performance is not significantly better than the 95^th^ percentile of models trained on permuted data. (B) On the left, correlation between the total cross-sectional area of early-visual tracts and surface-area of the posterior functional system, across subjects. On the right: a tractogram showing the specific hippocampal-cortical tracts belonging to the early-visual cortex.

We found that the size of the posterior system could be significantly predicted from the cross-sectional areas of early_visual, primary_visual, medial_temporal, posterior_cingulate, lateral_temporal, dorsal_stream_visual, and ventral_stream visual tracts; the size of the anterior functional system could be significantly predicted from primary_visual and medial_temporal tracts; while the size of the hippocampal body could be significantly predicted from posterior_cingulate tracts (Fig. 4A). Sensitivity analyses (see methods) indicated this effect was not dependent on sex or MRI acquisition site.

The best out-of-sample performance (out-of-sample *R^2^ =* .097, outperforming all 500 permutations) came from the model predicting the size of the posterior system from the cross-sectional areas of early-visual tracts. This association is highlighted in Fig. 2B, where we plot the sum of these early-visual tracts against the size of the posterior system (left scatter-plot; regression statistics:□ = 9.41 ± 2.08, *R*^2^ = .049, *p* = 7.6e-6) along with the 11 early-visual tracts contributing to this sum (right tractogram).

### Developmental changes in occipital fiber tracts contribute to development of the posterior functional system

To test our hypothesis that developmental changes in functional system surface-area may be partially attributable to developmental changes in the cross-sectional area of hippocampal fiber tracts, we used a simple mediation analysis (Baron & Kenny, 1986; Preacher & Hayes, 2004, 2008; MacKinnon et al., 2004). This involved fitting a set of three linear regressions estimating: (1) the effect of age on tract cross-sectional area [*a*: CSA-mm^2^ / year], (2) the conditional effect of tract cross-sectional area on functional system surface-area controlling for age [*b*: SA-mm^2^ / CSA-mm^2^], and (3) the total effect of age on functional system surface-area [*c*: SA-mm^2^ / year]. From these, the indirect effect of fiber-tract area on the relationship between age and surface-area is calculated by taking the product of *a* and *b,* with significance calculated by examining the 95% confidence interval of 1000 bootstrapped samples (all implemented in the call to *mediation_analysis*: https://pingouin-stats.org/). To test our hypothesis, we used a conservative joint significance test of all paths in the analysis. We note that, because all variables were measured cross-sectionally, this mediation analysis evaluates statistical compatibility with a developmental pathway, but does not establish causality.

The result of this statistical mediation analysis is presented in Fig. 5. System–cortex pairs which showed a significant mediating effect are additionally bolded. Similar to our models predicting functional system size from fiber-tract areas where the posterior systems showed the strongest effect, we found that only the size of the posterior system (not the anterior/body systems) showed a mediating effect. Specifically, the tracts which showed a significant mediating effect on the relation between age and posterior-system surface-area were: early_visual [*a*: –0.28 CSA-mm^2^ / year; *b*: 9.4 SA-mm^2^ / CSA-mm^2^; *a*b*: −1.16 SA-mm^2^ / year] and lateral_temporal cortex [*a*: –.05 CSA-mm^2^ / year; *b*: 14.66 SA-mm^2^ / CSA-mm^2^; *a*b*: –.64 SA-mm^2^ / year (2.98% of *c*)]. The total effect of age on surface-area for all models (*c*) was – 21.45 mm^2^ / year (Fig.. 5B, right). The most reliable effect of mediation (*p*-value from bootstrapped confidence interval) was for Early_Visual cortex, which is highlighted in Fig. 5b. Sensitivity analyses (see methods) indicated this effect was not dependent on sex or MRI acquisition site.

**Fig 5.**
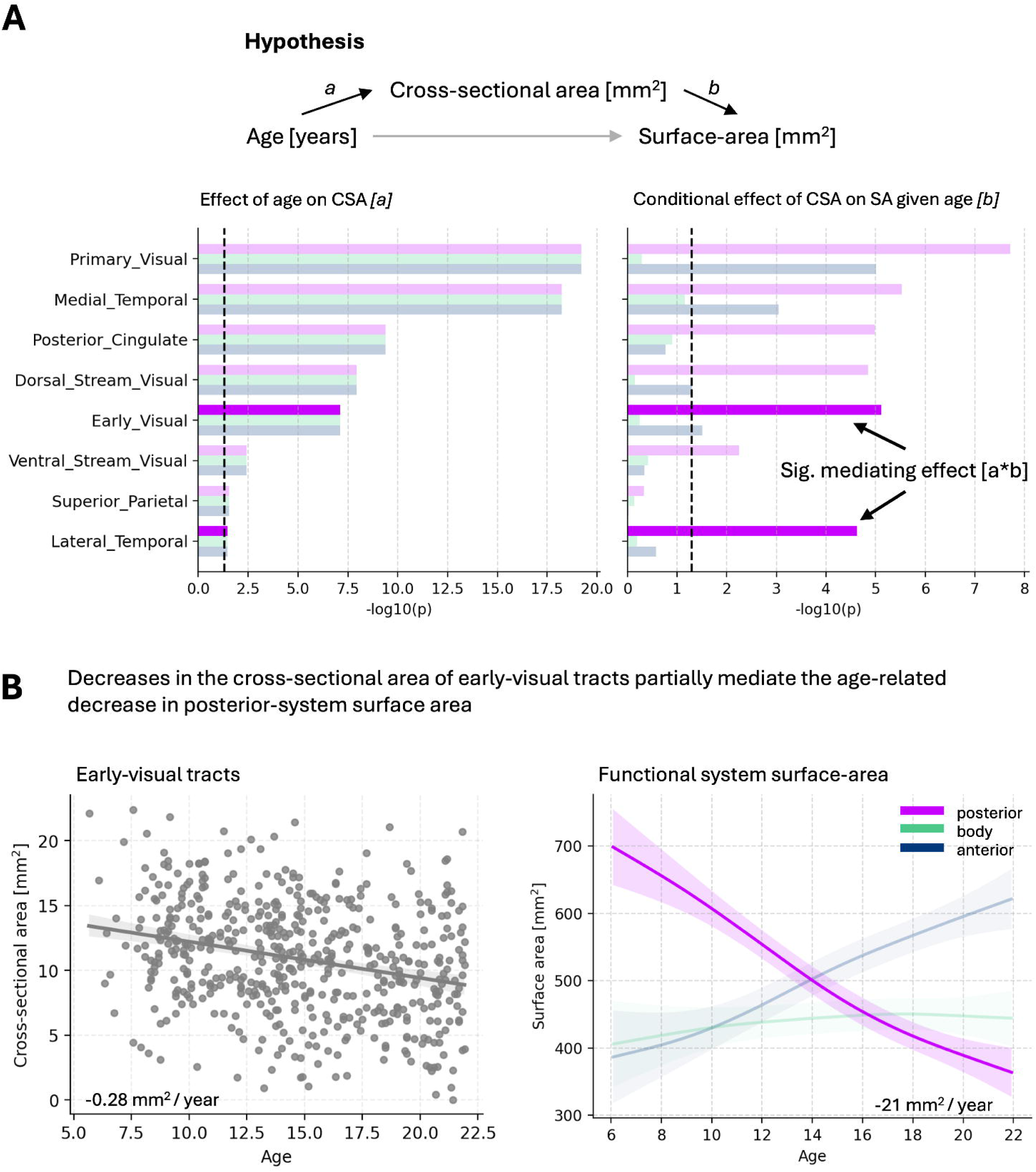
Developmental changes in occipital-parietal fiber-tracts contribute to development of the posterior functional system. (A) The hypothesized relation between variables, with fiber-tract cross-sectional area mediating the relation between age and functional-system surface-area. Bar-plots indicate *p*-values from the paths tested in the mediation analysis (dashed-line: *p =* .05). Effects with joint significance of paths (*a*, *b*, *a*b*) are bolded. (B) On the left, the correlation between age and cross-sectional area of early-visual tracts. On the right, age-related differences in the surface-area of each hippocampal system.

Together, these analyses provide correlational evidence consistent with the hypothesis that developmental pruning of the hippocampal fiber-tracts projecting to early-visual cortex (V2, V3, V4) contributes to developmental decreases in the size of the posterior functional system. Specifically, we found that early-visual fiber-tracts prune at a rate of approximately –.28 mm^2^ / year. Assuming linear effects, after controlling for age, each 1mm^2^ of pruning (equivalent to 3.57 years of development) is associated with a –4.1 mm^2^ decrease in surface-area of the posterior functional system. These parameter estimates suggest that each year of pruning results in a –1.16 mm^2^ decrease in posterior-system surface-area.

### Developmental changes in myelin content do not predict functional system surface-area after controlling for age

We conducted an identical set of analyses examining whether the T1w/T2w values of fiber-tracts could predict functional-system surface-area, and if so, whether fiber-tract T1w/T2w values had a mediating effect on the relation between age and functional-system surface-area.

Interestingly, we found that T1w/T2w of fiber-tracts was a reliable predictor of posterior system surface-area. This was true for fiber-tracts terminating in all cortices (see supplementary Fig. 2). For models trained to predict the anterior-system surface-area; lateral-temporal and ventral-stream-visual fiber-tracts also showed significant predictive performance. None of the models trained to predict the surface-area of the hippocampal body were significant.

Despite this predictive performance, simple linear regression models predicting surface-area from fiber-tract T1w/T2w while controlling for age, which were conducted as part of the mediation-analysis (path *b*), were not statistically significant (*p* < .05) for any cortex – functional-system pairs (see supplementary Fig. 2). Thus, while fiber-tract myelination is predictive of functional specialization in the hippocampus, this appears to be explained by the fact that both functional specialization and fiber-tract myelination increase with age.

## Discussion

We examined whether developmental differences in the strength of white-matter fiber-tracts could be a mechanism for functional specialization in the human hippocampus along the anterior–posterior axis. To do so, we non-invasively estimated the cross-sectional area of hippocampal fiber-tracts through a custom diffusion-MRI tractography pipeline, then demonstrated that these estimates can account for age-related increases in functional specialization. These findings provide correlational evidence in support of our hypothesized mechanism, whereby the anatomical development of hippocampal-cortical white-matter fiber-tracts over the course of childhood contributes to functional development of the hippocampus.

Specifically, we found that long-range fiber-tracts projecting between the posterior hippocampus and early-visual cortices (V2–V4; similar to those identified in Dalton et al., 2022) show small but reliable decreases in cross-sectional area throughout development. A probable mechanism for this effect is the stereotyped pruning of long-range axonal projections, a process which extends into adolescence (Bagri et al., 2003; Cressman et al., 2010; Riccomagno & Kolodkin, 2016). This developmental pruning was associated with decreased posterior-system surface-area, a sensitive marker of functional specialization in the hippocampus (Angeli et al., 2025; Kember et al., 2026). Statistical modeling further indicated a mediating effect of fiber-tract area on the relation between age and functional system surface-area. The direction of this effect is in line with the hypothesized mechanism. Indeed, the posterior hippocampal system is defined through correlated hemodynamic signaling with medial occipital-parietal cortex (Barnett et al., 2021; Zheng et al., 2021). In theory, by pruning the axons that connect the posterior hippocampus with occipital cortex, the synaptic efficacy of this connection is decreased, resulting in decreased neuronal and hemodynamic coactivation, and therefore a smaller functionally defined posterior-system.

We also observed a large increase in the cross-sectional area of short-range fiber-tracts terminating in the medial-temporal cortex. This process was significantly accelerated in the anterior compared to the posterior hippocampus, and was independent from the increases in myelin content observed within these tracts (e.g., not significantly correlated: *r*=-.02; in line with prior work correlating cross-sectional area with myelin content in short-range streamlines: see Nelson et al., 2023 supplementary). A possible explanation for this effect is an increase in the number of axonal projections, and/or an increase in the volume of the extracellular white-matter space (consisting of e.g., glia, interstitial fluid). Together, these increases in cross-sectional area and myelin content likely contribute to the rapid growth in white-matter volume seen during this time.

These findings have important implications for work aimed at promoting neurodevelopmental outcomes. Atypical functional development of the hippocampus is associated with deficits in learning, memory, and socioemotional regulation (Gilchrist et al., 2018). Our results suggest that atypical development of hippocampal-cortical white-matter tracts may contribute to altered functional specialization, highlighting these pathways as potential targets for future longitudinal and interventional studies.

## Limitations

These results are limited by: (1) the cross-sectional nature of our design, which limits the extent to which age-related differences can be interpreted as developmental effects. (2) The lack of specificity for T1w/T2w values to myelin content. (3) The directionality of the effect cannot be directly tested (i.e., while we infer that fiber-tract area influences functional specialization, the reverse could be true, with fiber-tract area refined through activity-dependent mechanisms).

## Methods

### Participants

All unrelated participants from the Human Connectome Project-Development cohort were used in this study (*N*=545, HCP-D; Somerville et al., 2018). All subjects were considered healthy, with no history of psychiatric illness (e.g., schizophrenia, anxiety, depression, etc.), head injuries, or the reception of support services in school (e.g., related to dyslexia, speech/language pathology, etc.). A full list of inclusion/exclusion criteria can be found in (Somerville et al., 2018). Informed consent was acquired during data collection. Four subjects were removed due to low-quality hippocampal segmentations (*HippUnfold* dice coefficients below 0.7; DeKraker et al., 2022); two subjects were removed due to low-quality hippocampal tractogram reconstructions (extreme outlying values in cross-sectional area), resulting in a total of 539 subjects entering analyses (aged 5-21 years, M±SD: 14.66 ± 4.18 years; 289 female, 250 male).

### MRI data acquisition

MRI data were collected on a Siemens 3T Prisma system equipped with 80 mT/m gradients (slew rate 200 T/m/s) using a 32-channel Prisma head coil. Both T1-weighted and T2-weighted images were acquired with 0.8 mm isotropic resolution (sagittal FOV = 256 × 240 × 166 mm; matrix = 320 × 300 with 208 slices), including 7.7% slice oversampling, two-fold in-plane acceleration along the phase-encoding direction, and a bandwidth of 744 Hz/pixel. For the T1-weighted image, sequence parameters were TR/TI = 2500/1000 ms, with four echo times (TE = 1.8, 3.6, 5.4, and 7.2 ms), a flip angle of 8°, and up to 30 repetitions. Water excitation was used to suppress fat signal from bone marrow and scalp tissue. For the T2-weighted image, TR/ TE = 3200/564 ms, turbo factor = 314, and up to 25 repetitions.

Diffusion-weighted images were acquired with 1.5 mm isotropic voxels using a multiband factor of 4, TR = 3.23 s, partial Fourier acquisition (6/8), and no in-plane acceleration. A total of 185 diffusion-encoding directions were sampled across two *b*-value shells (*b* = 1500 and 3000 s/mm²), with each direction acquired using both anterior–posterior and posterior–anterior phase encoding. Data were collected over four diffusion runs, with 28 interspersed *b* = 0 s/mm² volumes. Raw diffusion data were denoised using Marchenko–Pastur principal component analysis implemented in MRtrix3, followed by correction for head motion and eddy current–induced distortions using FSL, and correction for magnetic field nonlinearity using FreeSurfer.

### Hippocampal diffusion tractography

A volumetric segmentation of the hippocampus was obtained for each subject from their 0.8mm resolution T1-weighted image through *HippUnfold* (DeKraker et al., 2020). Following the hippocampal-tractography technique developed by Dalton et al. (2022), we modified these segmentations with an automated procedure whereby voxels 1mm below the inferior edge of the hippocampus were classified as ‘white-matter’ in the 5 tissue-type images used during streamline generation. This ensured that streamlines could innervate the inferior portion of the hippocampus in a biologically plausible manner, without modifying the location of the gray-matter white-matter interface itself.

Fiber orientation densities (FODs) were estimated in *MRtrix3* software using multi-shell multi-tissue constrained spherical deconvolution (CSD; *dwi2response dhollander*; *dwi2fod msmt_csd*; Tournier et al., 2019 [https://mrtrix.readthedocs.io/en/latest/]). These were then intensity normalized to allow for quantitative comparison of streamlines across subjects (measured in terms of cross-sectional area, mm^2^; done via the command *mtnormalise*; Raffelt et al., 2017). From these FODs and modified hippocampal masks, we generated 1 million anatomically constrained streamlines (500,000 from both the left and right hippocampi; relevant parameters: min-length: 3.0 mm; max-length: 150.0 mm; algorithm: iFOD2; -crop_at_gmwmi; - backtrack; -seed_unidirectional). To derive biologically accurate, quantitative estimates of fiber strength, we calculated the cross-sectional area of filtered streamlines by taking the sum of SIFT2 weights, scaled by the SIFT2 alpha parameter (Calamante et al., 2019; Dalton et al., 2022; Smith et al., 2015).

### Precision-mapping of hippocampal fiber-tracts

To summarize the set of hippocampal streamlines generated from our tractography pipeline, we first created a set of template hippocampal fiber-tracts, which captured common streamline trajectories across subjects. These templates were created via a two-step clustering procedure.

The first step of this procedure involved clustering all hippocampal streamlines (*N*=1,000,000) within each subject based on the similarity of their spatial trajectories. This was done with the QuickBundles streamline-clustering algorithm (Garyfallidis et al., 2012). QuickBundles clusters streamlines based on the “minimum average direct-flip distance” (MDF), which is simply the average Euclidean distance between a set of evenly-spaced points along streamline *i*, and a set of evenly-spaced points along streamline *j* (parameters: 12-point MDF; distance-threshold=5.0mm; the minimum of “flipped” directions is taken to account for the lack of an objective start-end versus finish-end within the algorithm). On average, this produced 3,685 ± 1,902 clusters per subject, resulting in 1,986,343 clusters in total. Next, the centroid of each cluster was transformed to MNI-space through the subject-specific non-linear transforms created as part of the Human Connectome Preprocessing pipeline (e.g., the transform ‘standard2acpc_dc’; Glasser et al., 2015).

The second step of this template-creation procedure involved clustering these MNI-space cluster centroids, which was also done via Quickbundles (12-point MDF, distance-threshold=15.0mm). This provided us with a total of 850 template tracts. The majority of these tracts were extremely local, terminating in spatially adjacent medial-temporal regions (including the gray-matter white-matter interface of the hippocampus itself), and accounted for a small amount of intra-axonal cross-sectional area. We therefore filtered out tracts that contained less than 0.2 mm^2^ of cross-sectional area on average across subjects, resulting in a total of *N*=107 tracts. Despite removing over 87% of streamline clusters, this filtering procedure maintained over 98% of the total streamline cross-sectional area across subjects. Each of the 107 template tracts was then manually inspected for quality by authors JK and XC. This resulted in the removal of two tracts, both of which took a biologically implausible path from the left posterior hippocampus to the thalamus via the corpus-callosum. This provided us with a final set of 105 tracts: 62 of which terminated in the medial-temporal cortex, and 43 of which terminated outside the medial-temporal cortex.

Once these group-level template fiber-tracts had been identified, we performed streamline template-matching at the subject level. Specifically, within each subject, the MDF distance of streamlines to the centroid of each template fiber-tract was calculated, and streamlines were assigned to the fiber-tract for which this distance was minimal. Then, to further ensure that each streamline showed a relatively similar trajectory to the group of streamlines that were assigned to a given fiber-tract, we performed an outlier-removal step. Specifically, the MDF distance of each streamline to its fiber-tract centroid was calculated (in subject native space), a log-normal distribution was fitted to these distances, and streamlines outside the 99^th^ percentile of this distribution were removed. Then, to ensure that this outlier-removal step did not impact estimates of cross-sectional area, we re-ran SIFT2 on the reduced set of cleaned streamlines. This step effectively redistributed the total cross-sectional area of all streamlines to the cleaned streamlines. This is shown in supplementary Fig 5.

This technique allowed us to keep streamlines in subject native-space, thereby maintaining individual variability, while also allowing us to reconcile specific fiber-tracts across subjects. Each tract was then assigned to one of the 22 major cortices in the Human Connectome multimodal parcellation (Glasser et al., 2016) based on the endpoint of the fiber-tract centroid. Statistics in the main text were calculated from these fiber-tracts, except for analyses focused on differences across the hippocampal long-axis. In these instances, streamlines were assigned to one of anterior, body, or posterior hippocampus based on the precise location of their hippocampal endpoint in the three-system template created in Kember et al. (2026). These templates approximately evenly divide the hippocampus into three parts along the longitudinal axis. Compared to the FreeSurfer probabilistic atlas of the hippocampus (Iglesias et al., 2015), the anterior hippocampal system has the highest spatial overlap with: molecular_layer_HP-head (dice=.83), CA1-head (dice=.74), and GC-ML-DG-head (.60). The body hippocampal system had the highest spatial overlap with: molecular_layer_HP-body (dice=.56), CA1-body (dice=.56), subiculum-body (dice=.48). The posterior hippocampal system had the highest spatial overlap with: Hippocampal_tail (dice=.90), subiculum-body (dice=.55), presubiculum-body (dice=.43).

Generalized additive models [https://pygam.readthedocs.io/] were used to visualize non-linear changes in fiber-tract properties (cross-sectional area, myelin content) as a smooth function of age (Serven et al., 2018).

### Functional system surface-area

The surface-areas of hippocampal systems for each subject were obtained from our prior analyses (see Kember et al., 2026 for further details). In brief, system surface-areas were calculated by: projecting the BOLD signal (2.0 mm isotropic voxel-resolution; 22 ± 4 minutes of low-motion data across subjects; framewise-displacement < 0.25mm) to the midthickness of the hippocampus; correlating this signal (Pearson’s *r*) with all vertices on the cortical surface (fsLR_32k); assigning each vertex of the hippocampus midthickness to a “template” cortical system with which it was most correlated; then summing the surface-areas of hippocampal vertices within each functionally defined system. The precise templates used in this technique can be seen in Kember et al. (2026).

### Predicting functional system surface-area from fiber-tract strength

To assess whether the strength of fiber-tracts (e.g., cross-sectional area, myelination) could predict the surface-areas of functional systems, we fitted cross-validated ridge-regression models in scikit-learn ([https://scikit-learn.org/stable/index.html]). This was done through a *Pipeline()* object, which included a *StandardScaler()* and a *RidgeCV()* estimator. L2 regularization of this estimator was tested on a set of 50 alpha values logarithmically spaced between 1e-3 and 1e5, with the optimal value selected through 5-fold cross-validation. Out-of-sample performances and null distributions for significance testing were obtained through *permutation_test_score*().

### Sensitivity analyses

We assessed the sensitivity of our primary results to sex (Male/Female) and/or MRI acquisition site (UCLA, Harvard, UMinn, WashU) by one-hot encoding these variables and regressing their effects from the measures used in analyses. We found that the regression models predicting posterior-system surface-area from fiber-tract cross-sectional areas showed significant out-of-sample performance when regressing out: sex (out-of-sample *R*^2^ = .094, *p* < .002), site (out-of-sample *R*^2^ = .097, *p* < .002), as well as sex and site together (out-of-sample *R*^2^ = .094, *p* < .002). This is shown in supplementary Fig 4. We also found a significant mediating effect of early-visual cross-sectional area on the relationship between age and posterior-system surface-area after regressing out the effect of: sex (Indirect-□*: −1.23, p* = .03), site (Indirect-□*: −1.16, p* = .03), as well as sex and site together (Indirect-□*: −1.24, p* = .02).

## Data availability

Data necessary to re-create the primary results, including subject-level measures of fiber-tract cross-sectional area, fiber-tract myelin content, and functional-system surface areas, are included in [supplementary_data.csv]. Raw data are openly available, with permissions, from: [https://www.humanconnectome.org/study/hcp-lifespan-development/data-releases]

## Code availability

All code necessary to re-create the primary analyses have been shared to GitHub: [https://github.com/JonahKember/HCPD_dMRI_analyses].

## Author contributions

**Jonah Kember:** Conceptualization, Formal analysis, Methodology, Data Curation, Writing - Original Draft, Writing - Review & Editing, Visualization, Funding acquisition.

**Christine L. Tardif:** Writing - Review & Editing, Conceptualization, Supervision.

**Sylvain Baillet:** Writing - Review & Editing, Supervision.

**Ying He:** Writing - Review & Editing.

**Sam Audrain**: Writing - Review & Editing.

**Alex Barnett:** Writing - Review & Editing.

**Tracy Riggins:** Writing - Review & Editing.

**Xiaoqian Chai:** Writing - Review & Editing, Conceptualization, Supervision, Funding acquisition.

## Notes

### Competing Interest Statement

The authors have declared no competing interest.

## References

Almeida, R. G., & Lyons, D. A. (2017). On myelinated axon plasticity and neuronal circuit formation and function. Journal of Neuroscience, 37(42), 10023–10034.

Angeli, P. A., DiNicola, L. M., Saadon-Grosman, N., Eldaief, M. C., & Buckner, R. L. (2025). Specialization of the human hippocampal long axis revisited. Proceedings of the National Academy of Sciences, 122(3), e2422083122.

Audrain, S., Milleville, S., Wilson, J., Baffoe-Bonnie, J., Gotts, S., & Martin, A. (2024). The development of functional connectivity along the hippocampal long-axis in infants.

Ayhan, F., Kulkarni, A., Berto, S., Sivaprakasam, K., Douglas, C., Lega, B. C., & Konopka, G. (2021). Resolving cellular and molecular diversity along the hippocampal anterior-to-posterior axis in humans. Neuron, 109(13), 2091–2105.e6. 10.1016/j.neuron.2021.05.003

Bagri, A., Cheng, H. J., Yaron, A., Pleasure, S. J., & Tessier-Lavigne, M. (2003). Stereotyped pruning of long hippocampal axon branches triggered by retraction inducers of the semaphorin family. Cell, 113(3), 285–299.

Bajada, C. J., Schreiber, J., & Caspers, S. (2019). Fiber length profiling: A novel approach to structural brain organization. Neuroimage, 186, 164–173.

Barch, D. M., Shirtcliff, E. A., Elsayed, N. M., Whalen, D., Gilbert, K., Vogel, A. C.,… & Luby, J. L. (2020). Testosterone and hippocampal trajectories mediate relationship of poverty to emotion dysregulation and depression. Proceedings of the National Academy of Sciences, 117(36), 22015–22023.

Barnett, A. J., Reilly, W., Dimsdale-Zucker, H. R., Mizrak, E., Reagh, Z., & Ranganath, C. (2021). Intrinsic connectivity reveals functionally distinct cortico-hippocampal networks in the human brain. PLoS biology, 19(6), e3001275.

Calamante, F. (2019). The seven deadly sins of measuring brain structural connectivity using diffusion MRI streamlines fibre-tracking. Diagnostics, 9(3), 115.

Cressman, V. L., Balaban, J., Steinfeld, S., Shemyakin, A., Graham, P., Parisot, N., & Moore, H. (2010). Prefrontal cortical inputs to the basal amygdala undergo pruning during late adolescence in the rat. Journal of Comparative Neurology, 518(14), 2693–2709.

Dalton, M. A., D’Souza, A., Lv, J., & Calamante, F. (2022). New insights into anatomical connectivity along the anterior–posterior axis of the human hippocampus using in vivo quantitative fibre tracking. elife, 11, e76143.

DeKraker, J., Lau, J. C., Ferko, K. M., Khan, A. R., & Köhler, S. (2020). Hippocampal subfields revealed through unfolding and unsupervised clustering of laminar and morphological features in 3D BigBrain. NeuroImage, 206, 116328.

DeKraker, Jordan, Roy AM Haast, Mohamed D. Yousif, Bradley Karat, Jonathan C. Lau, Stefan Köhler, and Ali R. Khan. “Automated hippocampal unfolding for morphometry and subfield segmentation with HippUnfold.” elife 11 (2022): e77945.

Garyfallidis, E., Brett, M., Correia, M. M., Williams, G. B., & Nimmo-Smith, I. (2012). Quickbundles, a method for tractography simplification. Frontiers in neuroscience, 6, 175.

Gilchrist, C., Cumberland, A., Walker, D., & Tolcos, M. (2018). Intrauterine growth restriction and development of the hippocampus: implications for learning and memory in children and adolescents. The Lancet Child & Adolescent Health, 2(10), 755–764.

Glasser, M. F., & Van Essen, D. C. (2011). Mapping human cortical areas in vivo based on myelin content as revealed by T1-and T2-weighted MRI. Journal of neuroscience, 31(32), 11597–11616.

Glasser, M. F., Sotiropoulos, S. N., Wilson, J. A., Coalson, T. S., Fischl, B., Andersson, J. L.,… & Wu-Minn HCP Consortium. (2013). The minimal preprocessing pipelines for the Human Connectome Project. Neuroimage, 80, 105–124.

Hagmann, P., Sporns, O., Madan, N., Cammoun, L., Pienaar, R., Wedeen, V. J.,… & Grant, P. E. (2010). White matter maturation reshapes structural connectivity in the late developing human brain. Proceedings of the National Academy of Sciences, 107(44), 19067–19072.

Huang, C. C., Rolls, E. T., Hsu, C. C. H., Feng, J., & Lin, C. P. (2021). Extensive cortical connectivity of the human hippocampal memory system: beyond the “what” and “where” dual stream model. Cerebral Cortex, 31(10), 4652–4669.

Iglesias, J. E., Augustinack, J. C., Nguyen, K., Player, C. M., Player, A., Wright, M.,… & Alzheimer’s Disease Neuroimaging Initiative. (2015). A computational atlas of the hippocampal formation using ex vivo, ultra-high resolution MRI: Application to adaptive segmentation of in vivo MRI. Neuroimage, 115, 117–137.

Insausti, R., & Munoz, M. (2001). Cortical projections of the non entorhinal hippocampal formation in the cynomolgus monkey (Macaca fascicularis). European Journal of Neuroscience, 14(3), 435–451.

Kjelstrup, K. B., Solstad, T., Brun, V. H., Hafting, T., Leutgeb, S., Witter, M. P., Moser, E. I., & Moser, M. B. (2008). Finite scale of spatial representation in the hippocampus. Science (New York, N.Y.), 321(5885), 140–143. 10.1126/science.1157086

Kember, J., He, Y., Gracia-Tabuenca, Z., Barnett, A., & Chai, X. (2026). The posterior hippocampus becomes topographically and functionally specialized with development.

Kember, J., Patenaude, P., Sweatman, H., Van Schaik, L., Tabuenca, Z., & Chai, X. J. (2024). Specialization of anterior and posterior hippocampal functional connectivity differs in autism. Autism Research, 17(6), 1126–1139.

Li, Y., Shen, M., Stockton, M. E., & Zhao, X. (2019). Hippocampal deficits in neurodevelopmental disorders. Neurobiology of learning and memory, 165, 106945.

Nelson, M. C., Royer, J., Lu, W. D., Leppert, I. R., Campbell, J. S., Schiavi, S.,… & Tardif, C. L. (2023). The human brain connectome weighted by the myelin content and total intra-axonal cross-sectional area of white matter tracts. Network Neuroscience, 7(4), 1363–1388.

Nichols, E. S., Blumenthal, A., Kuenzel, E., Skinner, J. K., & Duerden, E. G. (2023). Hippocampus long axis specialization throughout development: A meta analysis. Human Brain Mapping, 44(11), 4211–4224.

Nichols, E. S., Al-Saoud, S., Fang, M., Eagleson, R., de Vrijer, B., McKenzie, C.,… & Duerden, E. G. (2025). Early functional organization of the anterior and posterior hippocampus in the fetal brain. Cerebral Cortex, 35(12), bhaf327.

Pothuizen, H. H., Zhang, W. N., Jongen Rêlo, A. L., Feldon, J., & Yee, B. K. (2004). Dissociation of function between the dorsal and the ventral hippocampus in spatial learning abilities of the rat: a within subject, within task comparison of reference and working spatial memory. European Journal of Neuroscience, 19(3), 705–712.

Raffelt, D. A., Tournier, J. D., Smith, R. E., Vaughan, D. N., Jackson, G., Ridgway, G. R., & Connelly, A. (2017). Investigating white matter fibre density and morphology using fixel-based analysis. NeuroImage, 144(Pt A), 58–73. 10.1016/j.neuroimage.2016.09.029

Riccomagno, M. M., & Kolodkin, A. L. (2015). Sculpting neural circuits by axon and dendrite pruning. Annual review of cell and developmental biology, 31(1), 779–805.

Sandrone, S., Aiello, M., Cavaliere, C., Thiebaut de Schotten, M., Reimann, K., Troakes, C.,… & Dell’Acqua, F. (2023). Mapping myelin in white matter with T1-weighted/T2-weighted maps: discrepancy with histology and other myelin MRI measures. Brain Structure and Function, 228(2), 525–535.

Servén D., Brummitt C. (2018). pyGAM: Generalized Additive Models in Python. Zenodo. DOI: 10.5281/zenodo.1208723

Smith, R. E., Tournier, J. D., Calamante, F., & Connelly, A. (2015). SIFT2: Enabling dense quantitative assessment of brain white matter connectivity using streamlines tractography. NeuroImage, 119, 338–351. 10.1016/j.neuroimage.2015.06.092

Somerville, L. H., Bookheimer, S. Y., Buckner, R. L., Burgess, G. C., Curtiss, S. W., Dapretto, M.,… & Barch, D. M. (2018). The Lifespan Human Connectome Project in Development: A large-scale study of brain connectivity development in 5–21 year olds. Neuroimage, 183, 456–468.

Sotiropoulos, S. N., & Zalesky, A. (2019). Building connectomes using diffusion MRI: why, how and but. NMR in biomedicine, 32(4), e3752. 10.1002/nbm.3752

Strange, B. A., Witter, M. P., Lein, E. S., & Moser, E. I. (2014). Functional organization of the hippocampal longitudinal axis. Nature reviews. Neuroscience, 15(10), 655–669. 10.1038/nrn3785

Tang, L., Pruitt, P. J., Yu, Q., Homayouni, R., Daugherty, A. M., Damoiseaux, J. S., & Ofen, N. (2020). Differential functional connectivity in anterior and posterior hippocampus supporting the development of memory formation. Frontiers in Human Neuroscience, 14, 204.

Thompson, C. L., Pathak, S. D., Jeromin, A., Ng, L. L., MacPherson, C. R., Mortrud, M. T., Cusick, A., Riley, Z. L., Sunkin, S. M., Bernard, A., Puchalski, R. B., Gage, F. H., Jones, A. R., Bajic, V. B., Hawrylycz, M. J., & Lein, E. S. (2008). Genomic anatomy of the hippocampus. Neuron, 60(6), 1010–1021. 10.1016/j.neuron.2008.12.008

Tournier, R. E. Smith, D. Raffelt, R. Tabbara, T. Dhollander, M. Pietsch, D. Christiaens, B. Jeurissen, C.-H. Yeh, and A. Connelly. *MRtrix3*: A fast, flexible and open software framework for medical image processing and visualisation. NeuroImage, 202 (2019), pp. 116–37.

Vogel, J. W., La Joie, R., Grothe, M. J., Diaz-Papkovich, A., Doyle, A., Vachon-Presseau, E., Lepage, C., Vos de Wael, R., Thomas, R. A., Iturria-Medina, Y., Bernhardt, B., Rabinovici, G. D., & Evans, A. C. (2020). A molecular gradient along the longitudinal axis of the human hippocampus informs large-scale behavioral systems. Nature communications, 11(1), 960. 10.1038/s41467-020-14518-3

White, T.A., Miller, S.L., Sutherland, A.E. et al. Perinatal compromise affects development, form, and function of the hippocampus part one; clinical studies. Pediatr Res 95, 1698–1708 (2024). 10.1038/s41390-024-03105-7

Xie, H., Illapani, V. S. P., Reppert, L. T., You, X., Krishnamurthy, M., Bai, Y.,… & Sepeta, L. N. (2024). Longitudinal hippocampal axis in large-scale cortical systems underlying development and episodic memory. Proceedings of the National Academy of Sciences, 121(44), e2403015121.

Zheng, A., Montez, D. F., Marek, S., Gilmore, A. W., Newbold, D. J., Laumann, T. O., Kay, B. P., Seider, N. A., Van, A. N., Hampton, J. M., Alexopoulos, D., Schlaggar, B. L., Sylvester, C. M., Greene, D. J., Shimony, J. S., Nelson, S. M., Wig, G. S., Gratton, C., McDermott, K. B., Raichle, M. E.,… Dosenbach, N. U. F. (2021). Parallel hippocampal-parietal circuits for self- and goal-oriented processing. Proceedings of the National Academy of Sciences of the United States of America, 118(34), e2101743118. 10.1073/pnas.2101743118

