## Supplementary Material for "Fiber-tract development contributes to functional specialization in the human hippocampus"

**SFig 1. Full-brain tractogram results showing the concentration of cross-sectional in short-range streamlines across the cortex**

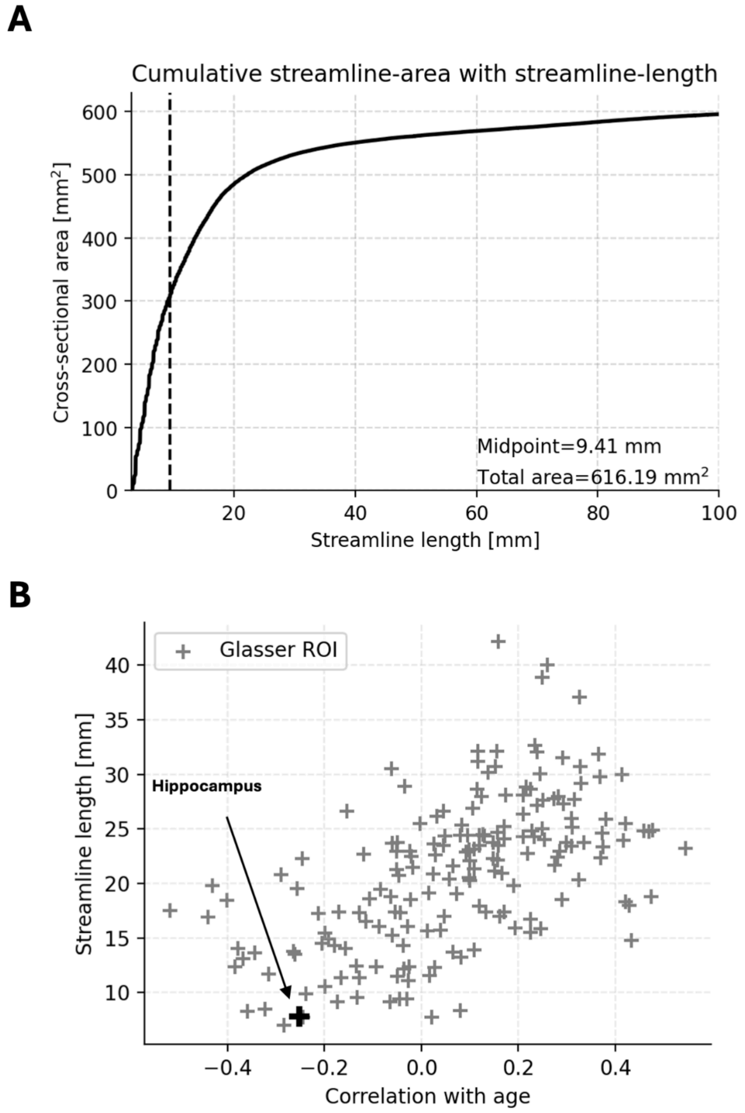


(A) Example of the process used to define the streamline-length that divided the total cross-sectional area of streamlines in two equal parts. This was used to measure the extent to which total streamline area was concentrated in short versus long streamlines.

(B) Scatterplot showing: streamline midpoints (as described in panel A) for all ROIs in the Glasser atlas, as well as the correlation between cross-sectional area and age or said ROI. As shown, the hippocampus is at an extreme end of the distribution, characterized by a high concentration of cross-sectional area in short-range streamlines. Moreover, a positive trend exists, where ROIs with a high concentration of area in short-range streamlines tend to decrease in area with age. The majority of ROIs show a positive correlation with age (116/180).

**SFig 2. Results predicting functional specialization from fiber-tract T1w/Tw2**

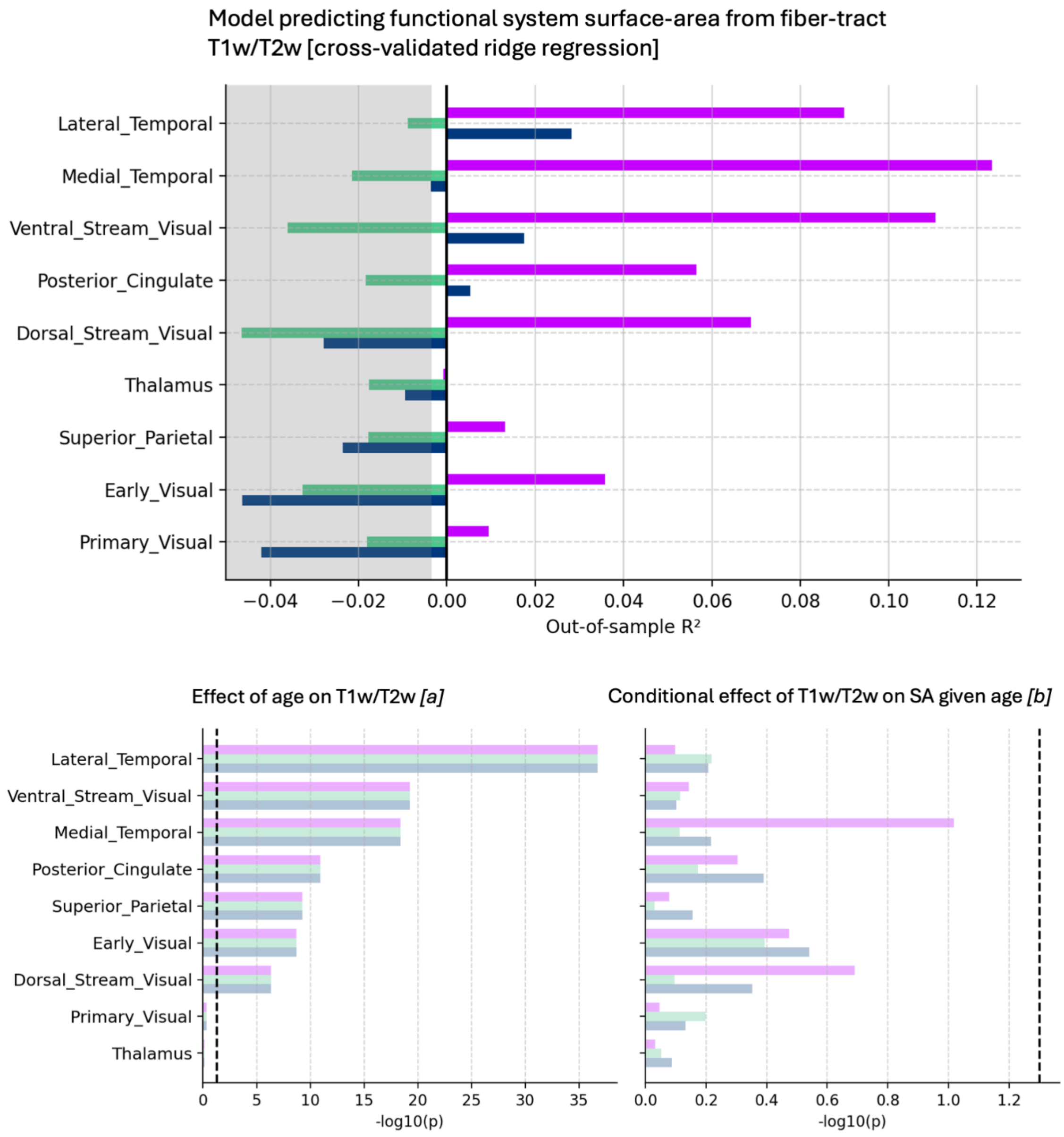

**SFig 3. Scatterplots of correlation between age and cortex-specific fiber-tract cross-sectional areas**

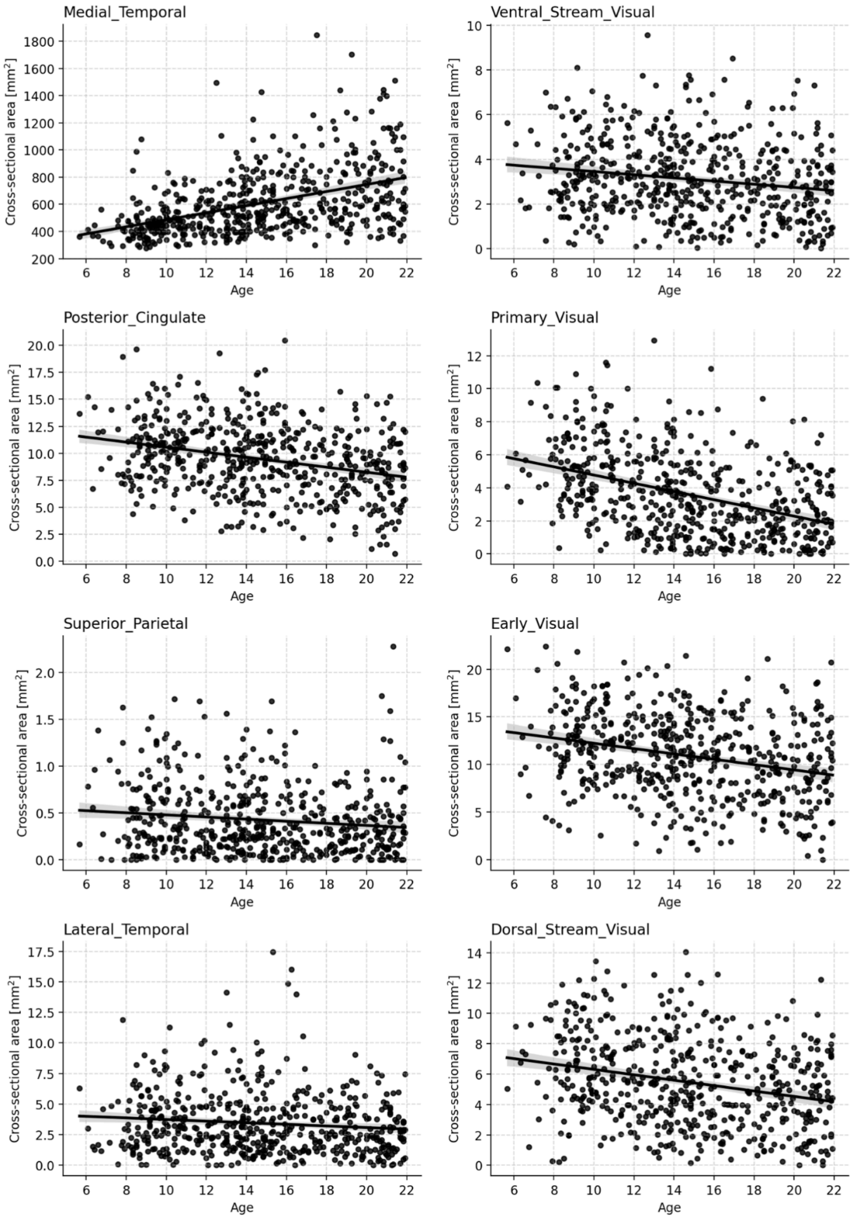


Correlations between fiber-tract cross-sectional area and age across subjects are shown.

**SFig 4. Sensitivity analyses of primary results adjusting for sex and MRI acquisition site**

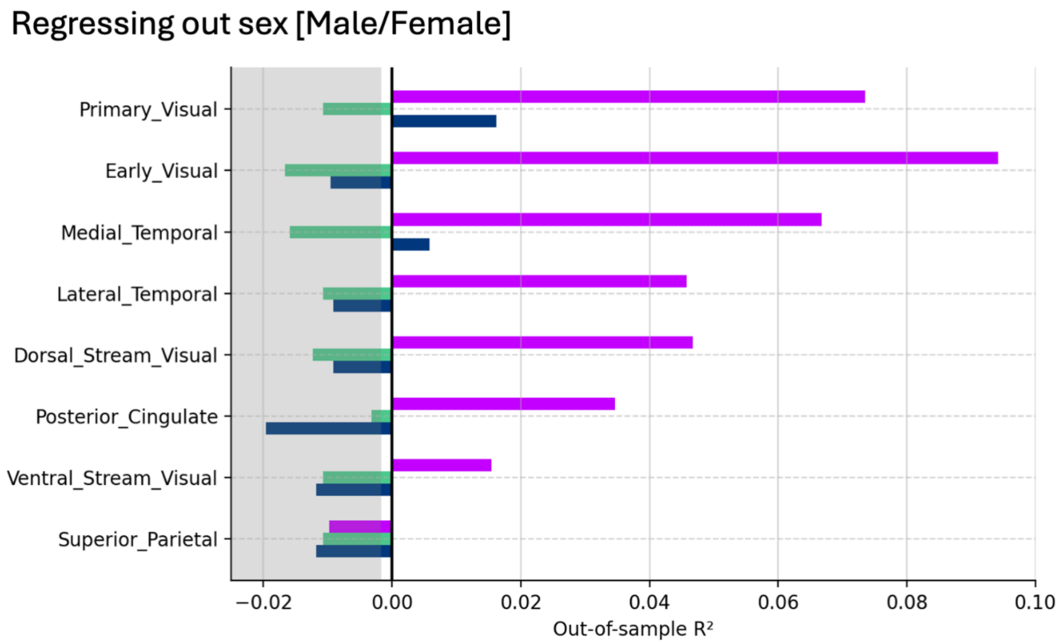


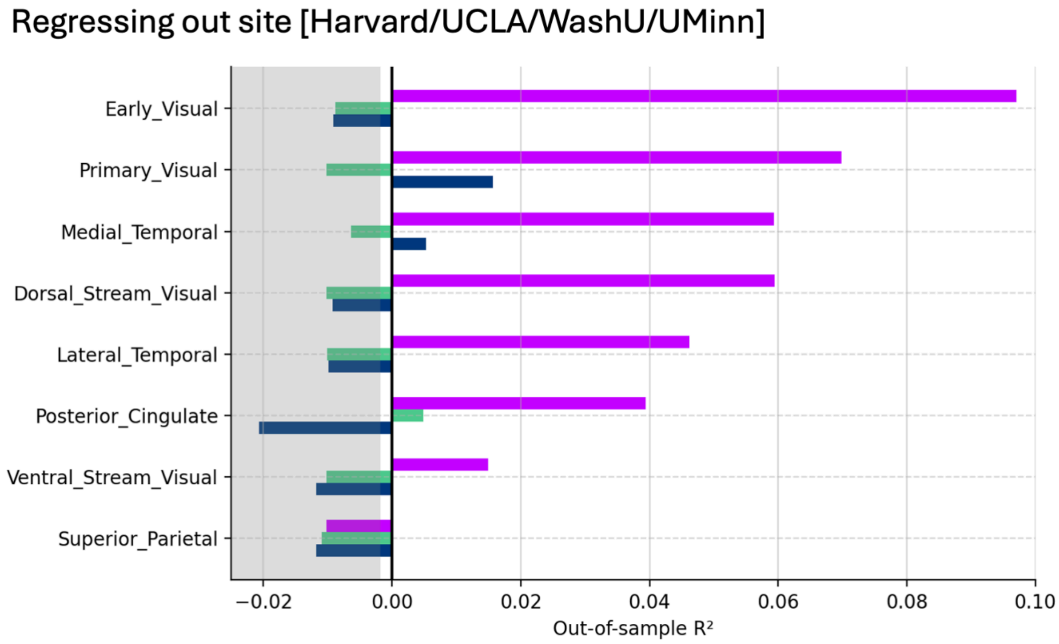


Results of penalized regression models predicting functional system surface-area from fiber-tract cross-sectional area are shown after regressing out sex and site.

**SFig 5. Streamline outlier removal detection**

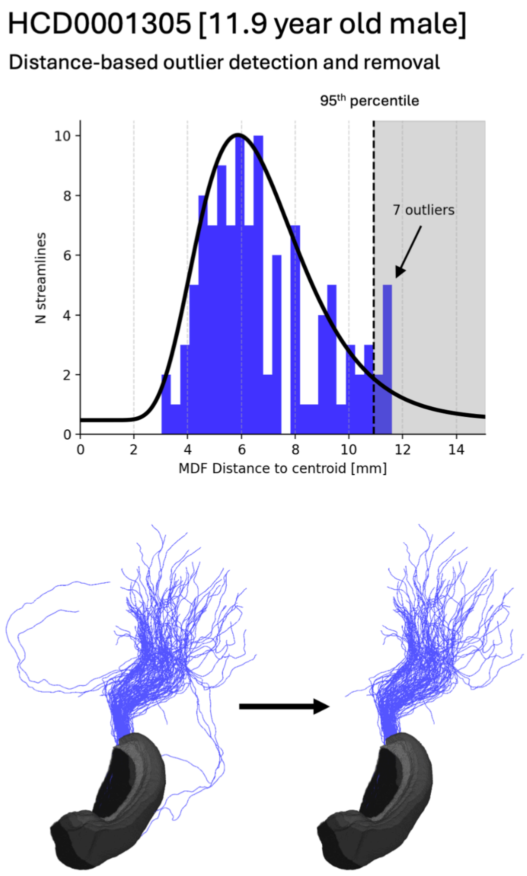


Example of the intermediate preprocessing step whereby streamlines assigned to a fiber-tract which show relatively high distance to the rest of streamlines assigned to said fiber-tract are filtered. SIFT2 is then run so as to re-distribute intra-axonal cross-sectional area to existing streamlines.

**SFig 6. Full hippocampus single subject tractogram**

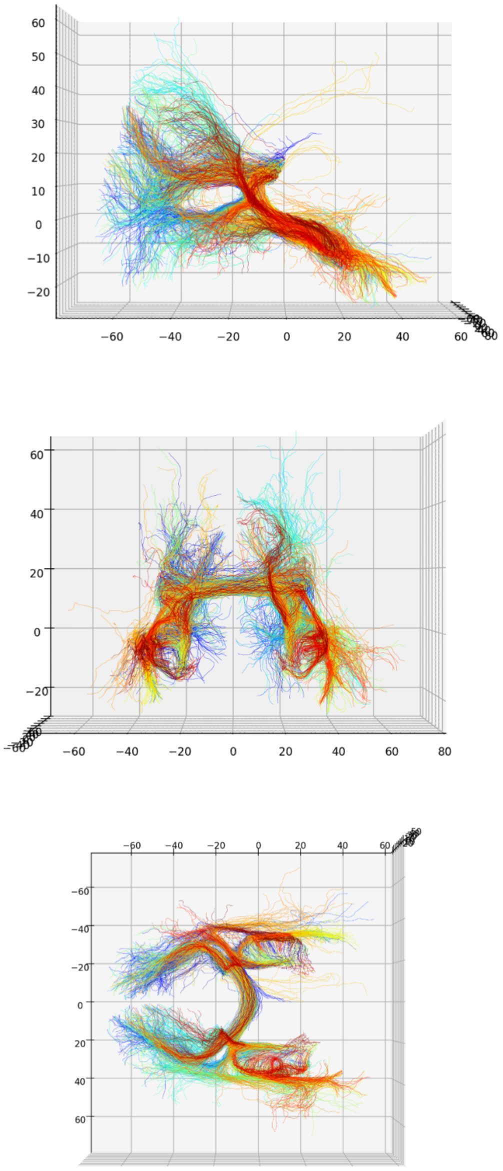


The hippocampal tractogram for a single subject. A subset of streamlines within each fiber-tract is shown to reduce computational demands

**SFig 7. Precision mapping of single-subject tractogram to template-tract**

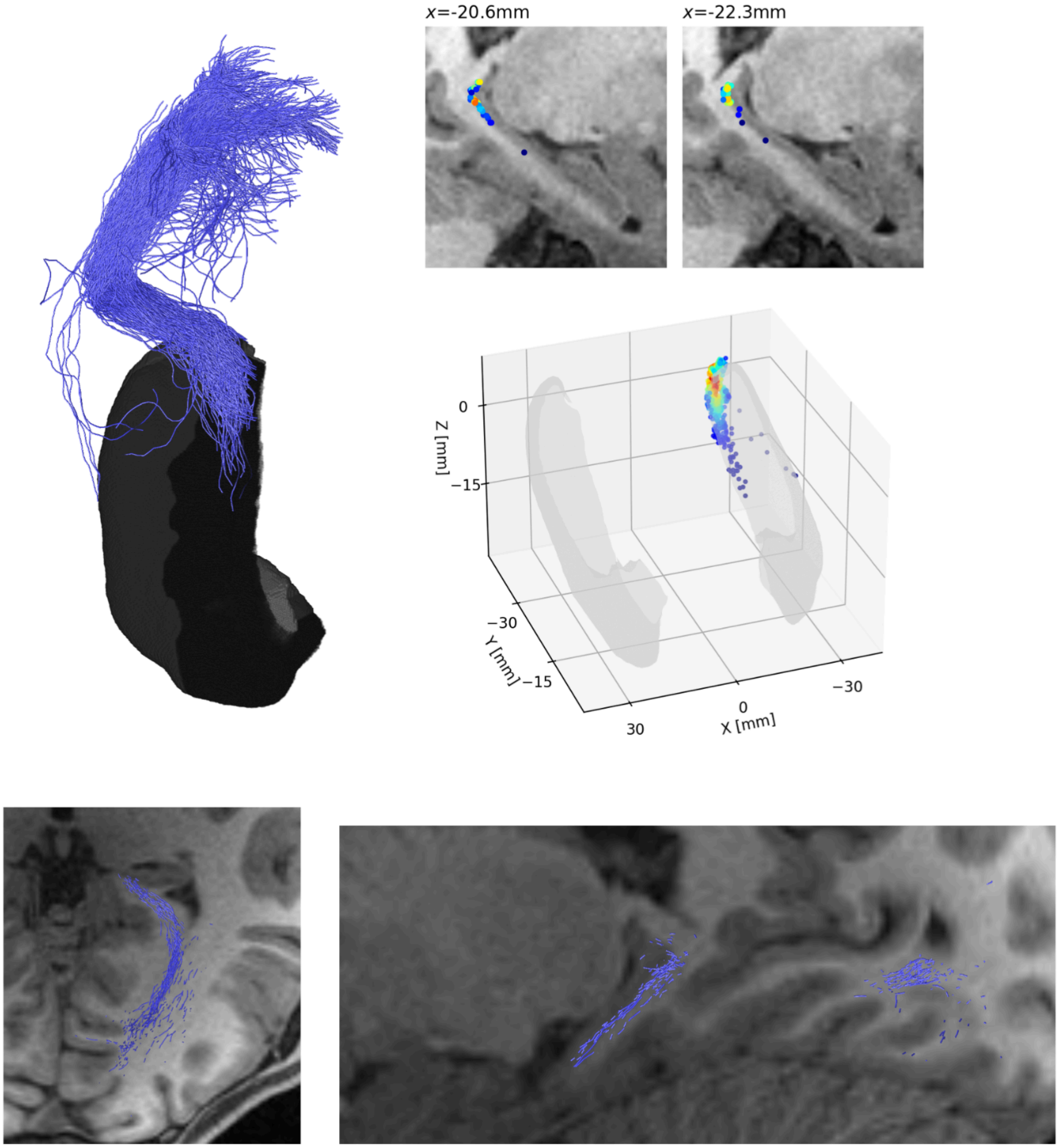


Streamlines are assigned to the template tract with which they showed the lowest MDF-distance. A collection of views for an example mapping of streamlines in a single subject to a single fiber-tract.
